# Comparative analysis of single-cell RNA sequencing methods with and without sample multiplexing

**DOI:** 10.1101/2023.06.28.546827

**Authors:** Yi Xie, Huimei Chen, Vasuki Ranjani Chellamuthu, Ahmad bin Mohamed Lajam, Salvatore Albani, Andrea Hsiu Ling Low, Enrico Petretto, Jacques Behmoaras

**Affiliations:** Programme in Cardiovascular and Metabolic Disorders, Duke-NUS Medical School, 8 College Road, 169857, Singapore, Singapore; Translational Immunology Institute, Singhealth/Duke-NUS Academic Medical Centre, the Academia, Singapore; Department of Rheumatology and Immunology, Singapore General Hospital, and SingHealth Duke-NUS Medicine Academic Clinical Programme, Duke-NUS Medical School, Singapore; Institute for Big Data and Artificial Intelligence in Medicine, School of Science, China Pharmaceutical University, Nanjing, 210009, China; Centre for Inflammatory Disease, Imperial College London, Hammersmith Hospital, London, W12 0NN

## Abstract

Single-cell RNA sequencing (scRNA-seq) has emerged as a powerful technique for investigating biological heterogeneity at the single-cell level in human systems and model organisms. Recent advances in scRNA-seq have enabled the pooling of cells from multiple samples into single libraries, thereby increasing sample throughput while reducing technical batch effects, library preparation time, and the overall cost. However, a comparative analysis of scRNA-seq methods with and without sample multiplexing is lacking. In this study, we benchmarked methods from two representative platforms: Parse Biosciences (Parse; with sample multiplexing) and 10X Genomics (10x; without sample multiplexing). By using peripheral blood mononuclear cells (PBMCs) obtained from two healthy individuals, we demonstrate that demultiplexed scRNA-seq data obtained from Parse showed similar cell type frequencies compared to 10X data where samples are not multiplexed. Despite a relatively lower library and cell capture efficiencies, Parse can detect rare cell types (e.g. plasmablasts and dendritic cells) which is likely due to its relatively higher sensitivity in gene detection. Moreover, comparative analysis of transcript quantification between the two platforms revealed platform-specific distributions of gene length and GC content. These results offer guidance for researchers in designing high-throughput scRNA-seq studies.

## Introduction

Single-cell RNA sequencing (scRNA-seq) has revolutionized the field of biomedical research by enabling the comprehensive profiling of mRNA expression levels at the single-cell level. This technology provides a means to unravel the inherent heterogeneity among cells, either through measuring proportional changes or alterations in gene expression [1]. Furthermore, recent advances in computational tools have facilitated the extraction of valuable insights from scRNA-seq data, including the exploration of cell differentiation trajectories and cell-cell communications [2, 3]. The exponential growth of scRNA-seq studies over the past decade has consolidated its wide usage as a powerful tool for addressing crucial questions in biomedical research. Nevertheless, the current limitations in sample-level throughput pose challenges for studies necessitating multiple conditions or samples, such as longitudinal studies. Traditional scRNA-seq protocols typically involve the preparation of a single sample in each library, which can inadvertently introduce batch effects when multiple samples are processed on different days and/or by different individuals. The confounding of batch effects with treatment conditions and the absence of biological replicates have been identified as significant factors contributing to false discoveries in scRNA-seq studies [4, 5]. Therefore, there is a critical need for methods that can increase sample throughput to ensure appropriately powered investigations [6].

One promising approach to address these challenges is the pooling of labeled samples into a mixture and parallelly profiling all cells with the aim of reducing technical batch effects, library preparation time, and overall cost. Various sample multiplexing techniques have been deployed to scRNA-seq protocols with different levels of sample throughput [7]. One of these, Parse Bioscience v2, commercialized the split-pool ligation-based transcriptome sequencing (SPLiT-seq) technology [8], enabling parallel profiling of 1-96 samples with up to 1 million cells in a single experiment. SPLiT-seq uses four rounds of combinatorial barcoding to index fixed and permeabilized cells parallelly without physically partitioning cells. The first round of barcoding serves the purpose of adding sample-level barcodes to cells. More specifically, each sample is distributed to a single well of a 96-well plate. Well-specific barcodes are appended to transcripts through an in-cell reverse transcription (RT) reaction. After this step, all the cells are pooled and redistributed to a new plate for the second round of barcoding. This split-pool-ligation procedure is repeated three rounds, resulting in a unique combination of three barcodes utilized as individual cell barcodes. Unique molecular identifiers (UMIs) are then added to each cDNA molecule at the third round to avoid amplification bias. After three rounds of barcoding, cells are split into 8 sublibraries where library specific barcodes are added and cDNA molecules are amplified. While commercially available plates from Parse Bioscience currently allow a maximum of 96 samples in one plate, the implementation of the first round of barcoding in a 384-well plate can easily scale up the system to accommodate 384 samples [8]. Despite the high-throughput potential of this platform, no study has yet compared its performance with conventional scRNA-seq protocols, where samples are not multiplexed, and transcriptomic profiles are generated independently for each sample.

Here, we systematically benchmarked Parse Bioscience v2 (hereinafter referred to as “Parse”) against conventional scRNA-seq protocol 3’ v3.1 (hereinafter referred to as “10x”) from 10x Genomics Chromium by comparing the transcriptomic profiles of peripheral blood mononuclear cells (PBMCs). The 10x platform utilizes a droplet-based scRNA-seq protocol, where each individual cell is captured and enclosed with a barcoded bead in a droplet using microfluidic system [9]. After reversion transcription within each droplet, droplets are merged by demulsification and cDNA amplification is done. We selected the 10x protocol for comparison because it is widely employed in scRNA-seq studies [10], and it has been extensively used to query human PBMCs for various questions such as new cell states common to healthy individuals [11] or specific to certain disease [12] or even existence of aging-related immune hallmark signatures [13]. For this platform comparison, we applied both Parse and 10x to the same PBMCs samples obtained from two ethnicity-matched healthy donors. The Parse replicates were multiplexed with nine other human PBMC samples, while the samples processed with the 10x protocol were not multiplexed, resulting in two independent libraries. We compared the transcriptomic profiles demultiplexed from Parse data with those obtained from a single 10x library. This allowed us to assess the library efficiency, gene detection sensitivity, accuracy in gene expression quantification and ability to recover biological distinct cell clusters between the two platforms.

## Results

### Study design and metrics

We benchmarked protocols (Parse *Evercode WT v2* and 10x *3’ v3.1*) from Parse and 10x platforms using PBMCs from two healthy donors (Fig. 1A, Supp. Figure 1). Human PBMCs serve as an ideal model system for benchmarking due to their heterogeneity in cell sizes and total RNA content, which are key factors affecting the performance of scRNA-seq methods [14]. Furthermore, human PBMCs are a well-studied cell system where the major cell types and their corresponding marker gene expression are well defined [11, 15-16]. Human PBMCs from two healthy donors (designated here as H1 and H2) were prepared in a single batch and distributed into two aliquots. Aliquot 1 was used for 10x library preparation without multiplexing H1 and H2 samples. Aliquot 2 was used for Parse library preparation by multiplexing H1 and H2 with nine other samples in a single library. All libraries were sequenced together to minimize differences in sequencing depth. For each platform, we aimed to collect ∼10,000 cells per sample.

**Fig. 1.**
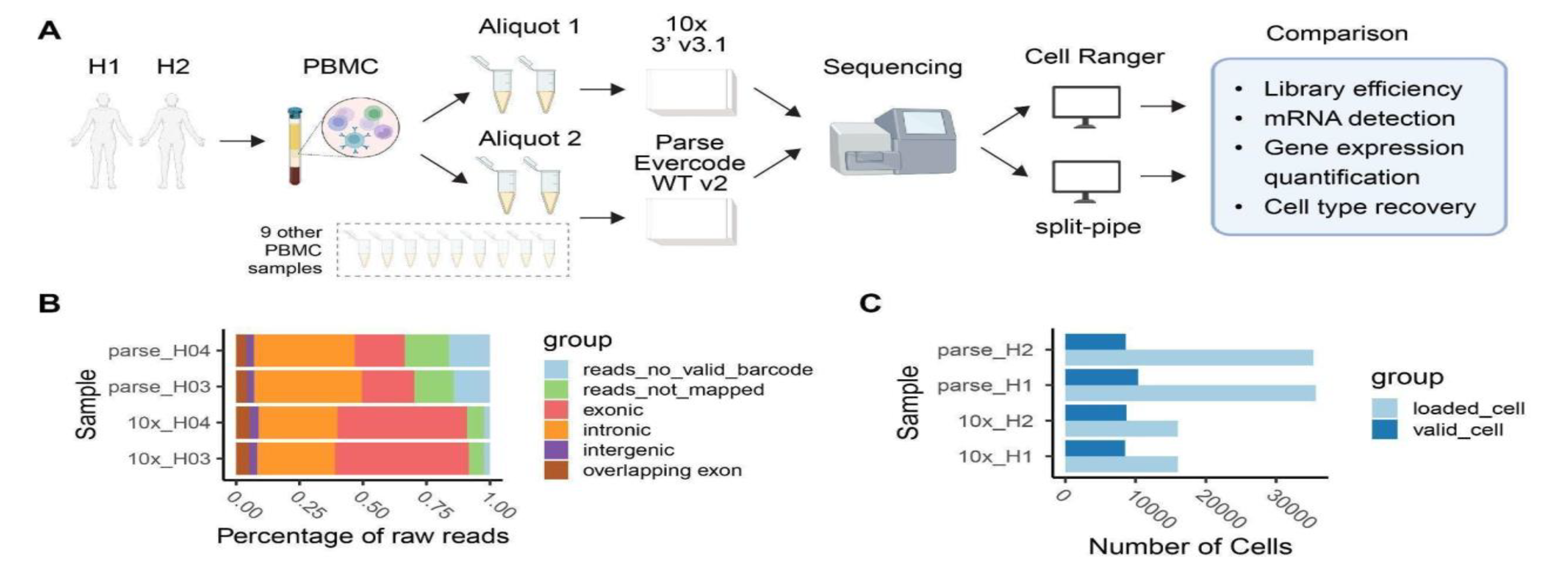
Overview of study design and library efficiency. **A)** Two human PBMC samples, H1 and H2, were split into two aliquots. One aliquot was used for 10x (3’ v3.1) library preparation and the other for Parse (Evercode WT v2) library preparation. These were sequenced parallelly on the same sequencer. Gene expression counts were generated separately, using pre-process pipelines specific to each method (*Cell Ranger* v7.1.0 [16] and *split-pipe* v1.0.4p). **B)** Percentage of raw reads discarded (reads_no_valid_barcode); not mapped to the genome (reads_not_mapped); mapped to exonic, intronic, intergenic and overlapping-exon regions. **C)** Number of cells loaded/input and number of valid cells output from pre-process pipelines.

In order to perform a comparative assessment of the two platforms, here we used four key metrics: library efficiency, gene detection sensitivity, accuracy in gene expression quantification and ability to recover distinct cell clusters. Library efficiency is directly related to the number of input cells and sequencing depth used in scRNA-seq studies [14]. As such, the two key factors that can affect library efficiency are cell recovery rate and fraction of reads with valid barcodes. Platforms that achieve a higher cell recovery rate are advantageous for studies that rely upon valuable (i.e., difficult to obtain) samples. Lower fraction of valid reads usually indicates high background noise [17] and requires deeper sequencing (and relatively increased cost) to ensure optimal transcriptome coverage. Sensitivity and accuracy in transcript detection and quantification will influence the ability to assess cellular heterogeneity. Higher sensitivity in gene detection will influence characterization of discrete cell clusters and transition to cell states, as well as cell-cell communications.

### Comparison of library efficiency

One factor affecting the scRNA-seq efficiency is the fraction of reads with valid cell barcodes. Too many invalid reads indicate the presence of high background noise in the library [14]. The fraction of valid reads was ∼ 85% for Parse and ∼ 98% for 10x (Fig. 1B). Valid reads were then mapped to the human genome to examine the distribution of reads throughout the genome. We observed that Parse had a higher proportion of intronic reads and lower proportion of exonic reads compared to 10x (Fig. 1B). One possible explanation for this result is that oligo-dT primers used in the 10x platform induced a biased priming towards exon regions next to 3’ polyA tails [18]. Parse reduced this bias by utilizing a mixture of oligo-dT and random hexamer primers [19].

Another significant factor in scRNA-seq efficiency metrics is the fraction of cells recovered in the data relative to input, namely cell capture efficiency. This factor is particularly important when one considers preparing libraries from biospecimens presenting a low number of cells. The number of valid cells recovered in each sample was directly obtained from 10x and Parse’s preprocessing pipelines. Both methods use barcode ranking plots to identify barcodes with valid cells and barcodes deriving from ambient RNA. We observed that on average ∼53% of cells were recovered in 10x, and ∼27% of cells were recovered in Parse data (Fig. 1C, Supp. Table 1).

**Table 1.** Summary of the comparative analysis between Parse Biosciences and 10x Genomics

| PLATFORM |  | 10x | Parse |
| --- | --- | --- | --- |
| STRENGTHS |  | - higher library efficiency | - enables tagging of 96 samples parallelly in a single batch with up to 1M cells<br>- higher sensitivity in gene detection |
| WEAKNESSES |  | - enables tagging of 16 (max) samples sequentially in a single lane with up to 10K cells<br>- lower sensitivity in gene detection | - significantly lower cell recovery rate |
| ASSESSED FEATURE | METRICS | 10x | Parse |
| LIBRARY EFFICIENCY | *cell recovery (average %)* | 53% | 27% |
|  | *valid reads (average %)* | 98% | 85% |
| SENSITIVITY IN GENE DETECTION | *number of genes (median [range])* | 1,938<br>[515 - 4,967] | 2,302<br>[860 - 5,548] |
| POTENTIAL BIASES IN GENE QUANTIFICATION | *gene length (median [range])* | 881 bp<br>[255 - 18,662] | 2,394 bp<br>[331 - 36,798] |
|  | *GC content (median [range])* | 49.4%<br>[36.8% - 71.6%] | 39.9%<br>[34.1% - 63.6%] |
|  | *ribosomal protein coding genes abundance (median [range])* | 30.7 %<br>[0.4% - 66.0%] | 1.0 %<br>[0% - 5.3%] |
| ABILITY TO IDENTIFY CELL TYPES ACCURATELY | *manual annotation<sup>1</sup> (average expression<sup>2</sup>, % of cells expressed<sup>3</sup>)* | B cells: 8.0, 93.9%<br>Monocytes: 102.0, 99.5%<br>NK cells: 26.8, 91.1%<br>T cells: 3.4, 80.3% | B cells: 4.2, 78.0%<br>Monocytes: 14.8, 74.9%<br>NK cells: 2.5, 58.7%<br>T cells: 0.8, 30.4% |
|  | *reference-based scoring<sup>4</sup> (median AUC [range])* | B cells: 0.12 [0.04 - 0.18]<br>Monocytes: 0.18 [0.12- 0.22]<br>NK cells: 0.11 [0.04 - 0.16]<br>T cells: 0.07 [0.02 - 0.11] | B cells: 0.14 [0.08 - 0.24]<br>Monocytes: 0.15 [0.02 - 0.19]<br>NK cells: 0.11 [0.04 - 0.16]<br>T cells: 0.08 [0.02 - 0.12] |

<sup>1</sup>we annotated each cell cluster using the expression of known cell type marker genes (see Methods).
<sup>2</sup>for each cell type, we report the the average expression of cell type markers across all cells in the cluster.
<sup>3</sup>percentage of cells in the cell type cluster that express the cell type markers.
<sup>4</sup>for each cell type cluster, we calculated a cell type activity score (AUC) using the gene signatures from reference dataset of sorted immune cells [27] (see Methods)

### Sensitivity in gene detection

To compare the sensitivity of mRNA capture between the two methods, we downsampled the sequencing reads to 20,000 per cell for each sample from 10x and Parse. Overall, we observed a ∼1.2 fold increase in the number of detected genes in Parse, with median value of 1,886 and 1,984 genes in 10x samples H1 and H2, respectively, compared to 2,319 and 2,283 genes in Parse samples H1 and H2, respectively (Fig. 2A, Supp. Table 2). Note that within each platform, the two biological replicates show a similar number of detected genes/UMIs, indicating high technical reproducibility for both 10x and Parse (Fig. 2A, Supp. Table 2-3). To better characterize the library complexity of two methods, we quantified the number of detected genes/UMIs at varying sequencing depths. We observed that both platforms have a similar number of detected UMIs while Parse consistently provides more genes detected at varying sequencing depths (Fig. 2B).

**Fig. 2.**
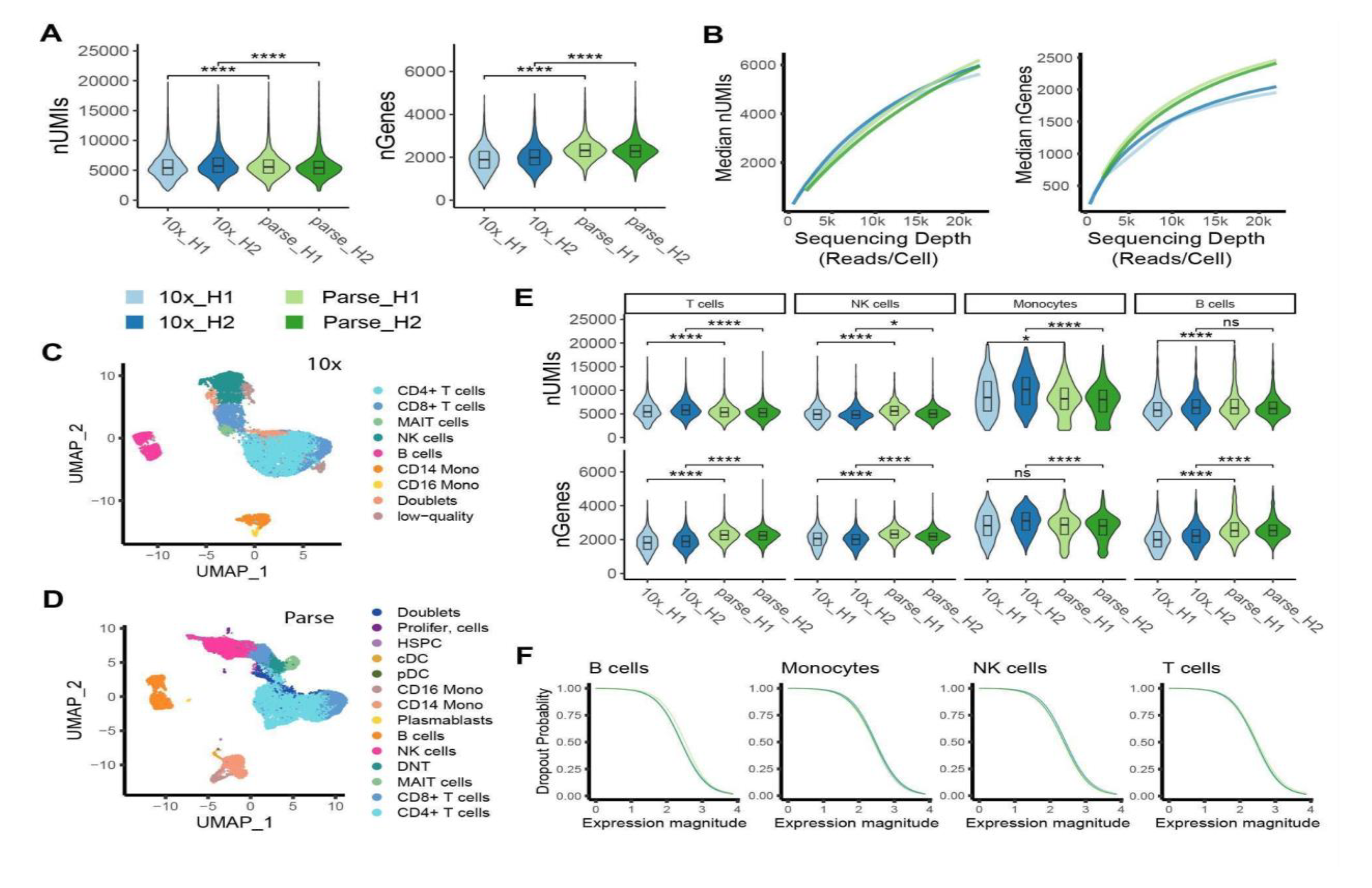
Sensitivity in capturing mRNA molecules. **A)** Median number of detected transcripts (UMIs) and genes (nGenes) at different down-sampled sequencing depth in each sample. **B)** Violin plot displaying the distribution of number of UMIs (left panel) and genes (right panel) per cell across all samples at sequencing depth 20,000 reads/cell. **C, D)** UMAP visualization of cell type annotations in 10x and Parse data. **E)** Violin plot displaying the distribution of number of UMIs (top) genes (bottom) per cell for the major cell types across all samples at sequencing depth 20,000 reads/cell. (ns: p > 0.05, *p <= 0.05, **p <= 0.01, ***p <= 0.001, two-samples *t*-test) **F)** Dropout probabilities plotted as a function of gene expression magnitude for each method (green and blue lines, respectively) for 4 major representative cell types in PBMCS.

After removing low-quality cells (cells with low UMI/gene counts or high percentage of mitochondrial mRNA) and doublets, we recovered 6,920 and 7,020 cells for samples H1 and H2, respectively, in 10x data. Similarly, the Parse dataset yielded 9,819 and 8,259 cells for samples H1 and H2, respectively (Supp. Table 1, Supp. Figure 2). Cell type identities were obtained by consensus annotation using cell type markers and reference datasets (see Methods; Fig. 2C, 2D; Supp. Figure 3). Sensitivity of mRNA capture was determined separately for major cell types in human PBMCs, namely T cells, NK cells, monocytes, and B cells. Across all cell types, Parse consistently exhibited the detection of a higher number of genes, with the exception of monocytes where the trend was reversed (Fig. 2E). Specifically, within the monocyte population, the 10x dataset detected an additional 187 genes compared to Parse (Supp. Table 2). This discrepancy is likely attributed to the improved performance of the 10x protocol in cells with higher RNA quantities, as it was previously shown [20].

**Fig 3.**
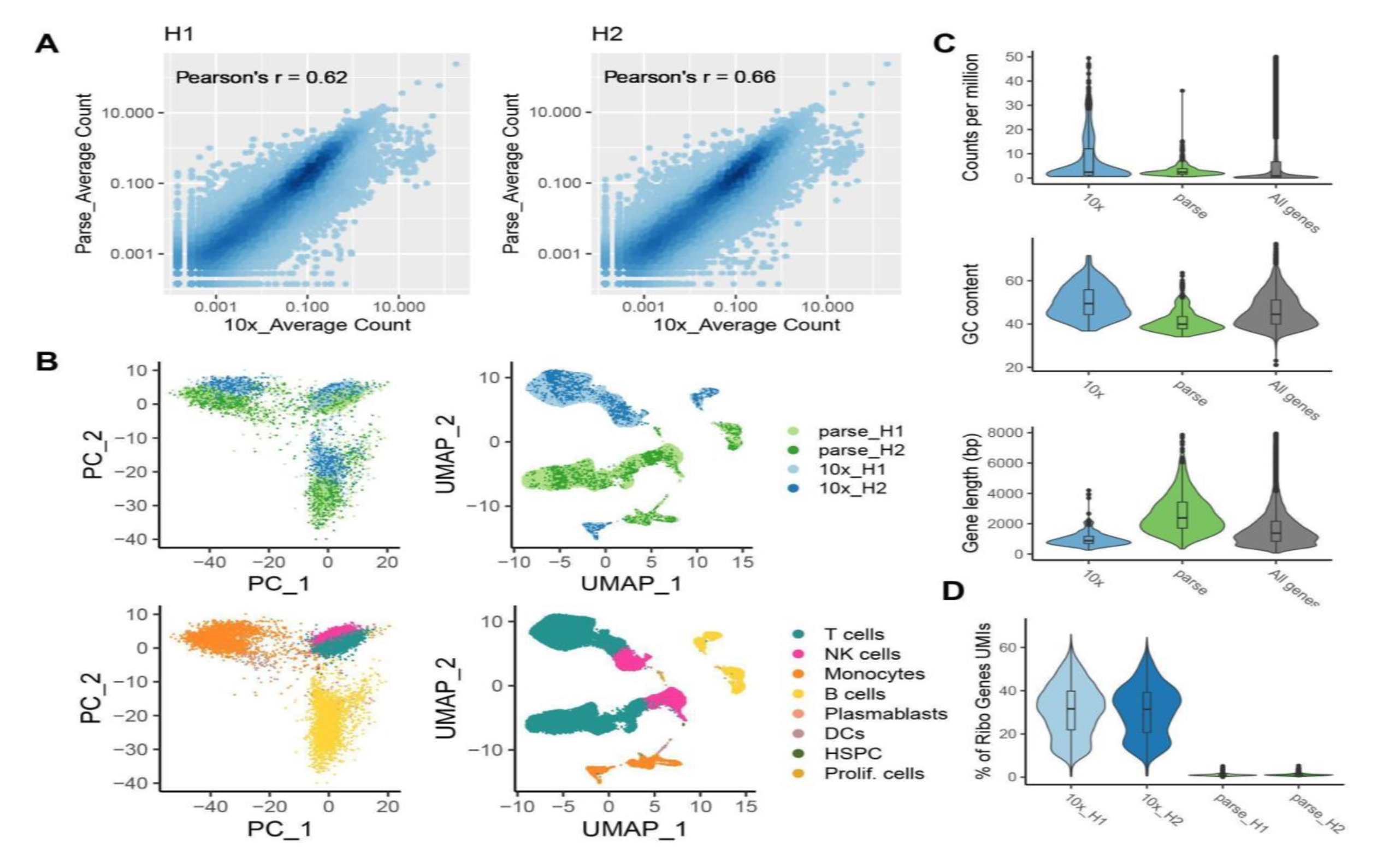
Comparative analysis of gene quantification bias. **A)** Density scatter plots showing gene expression correlation between the two platforms in sample H1 (left) and sample H2 (right). **B)** PCA and UMAP visualization of cells colored by methods and cell types. **C)** Distributions of gene expression abundance, gene GC content, gene length. Marker genes from each platform were used to demonstrate the distributions in the first two violins (317 genes for 10x; 556 genes for Parse). The distribution of all genes (29,184) is shown for comparison in the third violin. **D)** Percentage of UMIs mapped to ribosomal protein coding genes in each cell.

A dropout event is the phenomenon whereby lowly-expressed genes quantified in one cell are not detected in another cell, leading to challenges in downstream analysis such as differential expression analysis [21]. Here we estimated the ability of each platform to detect genes at various expression levels by calculating dropout probabilities. We observed that 10x and Parse had similar power to detect genes at varying expression levels (Fig. 2F).

### Bias in gene quantification

Genes expressed in less than 8 cells across all samples were considered lowly-expressed and removed from our dataset. In sample H1, there are 25,822 commonly expressed genes between two platforms, 2,035 Parse-specific genes and 1,702 10x-specific genes (Supp. Figure 4A). Similarly, for sample H2, we have 1,814 genes uniquely detected in Parse, 1,739 genes uniquely detected in 10x and 26,006 genes both detected by two platforms (Supp. Figure 4B).

**Fig 4.**
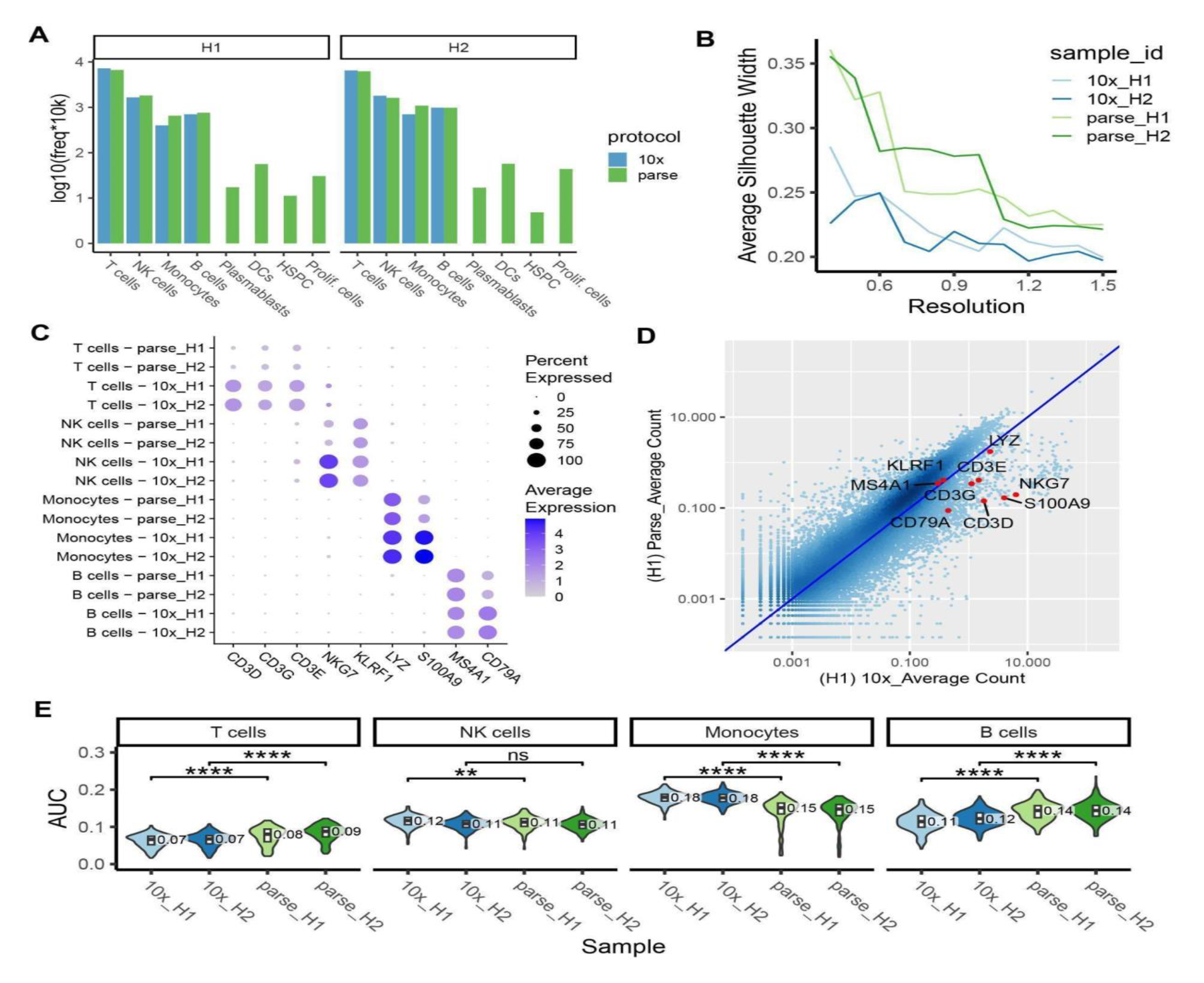
Clustering and cell type annotation. **A)** Barplots showing cell types identified in each method separately and the number of cells in each cell type. **B)** Clustering quality at different clustering resolutions measured by average silhouette width (ASW) score. **C)** Dotplot showing the expression of key cell type markers in two methods. The size of the dot encodes the percentage of cells expressing that marker and the color encodes the average expression level. **D)** Density scatterplot showing gene expression correlation between the two platforms in sample H1. X-axis is the average UMI count/gene in 10x data and Y-axis is the average UMI count/gene in Parse data. Cell type markers are highlighted as red dots. **E)** Cell type activity score calculated by *AUCell* for each cell type in each sample (ns: p > 0.05, *p <= 0.05, **p <= 0.01, ***p <= 0.001, two-samples *t*-test).

We then assessed the similarity of gene expression profiles detected by the two methods using correlation analysis. We randomly kept 7,000 cells for each sample in our downsampled data (20,000 reads/cell). 24,599 genes that are commonly expressed in all samples were used for correlation analysis. For each technical replicate, Pearson’s correlation coefficient (r) was calculated between two platforms based on the average UMI counts across all cells (Fig. 3A). We observed moderate variations between the two platforms with Pearson’s r = 0.62 for H1 and 0.66 for H2. These values are similar to previously reported comparisons between 10x and other platforms where Pearson’s r ranged from 0.5 to 0.7 [20]. Based on these findings, we conclude that the expression of genes is largely conserved across the two platforms.

To characterize variance between the transcriptomes depicted by two platforms, we performed dimension reduction with principal component analysis (PCA) and Uniform Manifold Approximation and Projection (UMAP). We observed that cells formed distinct cell type clusters including T cells, NK cells, B cells and Monocytes. Within each cell type cluster, cells are further clustered by their platform of origin, indicating technical differences existing between two platforms in gene quantification (Fig. 3B). Potential biases in gene quantification could be related to three major factors: gene expression abundance, gene length, GC content and PCR amplification bias [20, 22-23]. We examined the influence of these factors by selecting marker genes for each method and compared the distribution of these factors between two methods. We found that 10x has higher sensitivity to detect shorter genes and genes with higher GC content, which is consistent with previous findings [18], while Parse has higher sensitivity to detect longer genes and genes with lower GC content (Fig. 3C). Furthermore, our observations indicate that both the 10x and Parse platforms exhibited a comparable quantity of ribosomal protein coding genes (RB genes), with 424 and 425 genes detected in two samples of the 10x data; 316 and 319 genes detected two samples of the Parse data. However, the expression levels of these genes differ significantly between the two platforms. Specifically, the proportion of UMIs mapped to ribosomal protein coding genes varies from 0.4% to 66.0% in 10x, with a median value of 30.7%. In Parse, the range is from 0% to 5.3%, with a median value of 1.0% (Fig. 3D). We have also compared other scRNA-seq platforms, showing a range of enrichment for ribosomal protein coding genes in PBMCs (Supp. Figure 4G).

### Ability to distinguish and recover cell types

Identifying distinct cell types within a heterogeneous population is a fundamental step in analyzing scRNA-seq data. It is essential for identifying rare cell types, transitional cell states, and functionally significant cell subpopulations. Additionally, it facilitates advanced analysis such as differential gene expression between conditions and the investigation of cell-cell communication. Cell type identification is typically accomplished using two main methods: manual annotation and reference-based scoring. Our cell type annotation is achieved by harmonizing results from these two methods (see Methods). We observed that major cell types, T cells, B cell, NK cells and monocytes were identified by both platforms with consistent proportions (Fig. 4A), which are also aligned with cell type proportions previously reported in other scRNA-seq studies of human PBMCs [24, 25]. Rarer cell types (abundance < 1%), such as dendritic cells (DCs), plasmablasts, hematopoietic stem and progenitor cells (HSPC), and proliferating cells were identified by Parse only, which is consistent with the higher gene capture sensitivity results in improved cell cluster separation (Fig. 4A).

To further illustrate the ability of these two platforms to chart cell type heterogeneity, we utilized different metrics to quantify the power to distinguish and recover cell types with manual annotation or reference-based scoring. For the manual annotation method, we used average silhouette width (ASW) score and compared the expression of marker genes to compare the performance of 10x and Parse platforms. For the reference-based method, area under the receiver operating characteristic curve (AUC) was applied to quantify the platforms’ relative power to distinguish and recover cell types. These metrics enable us to quantitatively assess the effectiveness of the platforms in capturing and characterizing cell type diversity.

During manual annotation, cells with similar transcriptomic profiles are grouped into clusters based on known marker genes that are specifically and highly expressed in particular cell types. The clustering quality of two datasets was evaluated by calculating the average silhouette width (ASW) score, a commonly used measure for assessing data partitioning quality [26]. Consistent with its higher gene capture sensitivity, the Parse platform exhibited higher ASW scores at various resolutions in both samples (Fig. 4B). This finding was supported by Parse’s ability to effectively identify rare cell types (Fig. 4A). However, the expression levels of some cell type marker genes were significantly lower in the Parse dataset when compared to 10x (Fig. 4C). When analyzed by Parse, marker genes like *CD3D*, *CD3G*, *CD3E*, and *NKG7* were found to be expressed in less than 50% of cells within their respective cell types. Gene expression correlation analysis between the two platforms indicated that these specific cell type markers were detected in relatively more abundance by the 10x dataset (Fig. 4D, Supp. Figure 5), which compromises the Parse dataset’s ability to accurately annotate cell types using these specific markers.

We further assessed the ability of two platforms to recover cell types by reference-based scoring. Cell type specific gene signatures were derived from a reference dataset consisting of bulk RNA-seq samples generated from sorted and purified immune cell populations [27]. The area under the receiver operating characteristic curve (AUC) is calculated for each cluster to estimate how well cells in that cluster can be distinguished from the remaining cells (See Methods). The AUC summarized the performance of each cell type specific gene signature in separating a cluster of cells from the rest of the cells, with higher AUC values indicating better separability. We observed Parse exhibited higher AUC values in T cells, B cells and lower AUC values in monocytes (Fig. 4E). This observation is aligned with the trend of gene detection sensitivity in the two platforms, where a higher number of detected genes contributes to higher accuracy in identifying cell types.

## Discussion

We conducted a comprehensive comparison between the widely used 10x platform and Parse Biosciences, a relatively newly developed scRNA-seq platform that utilizes sample multiplexing and parallel profiling of a large number of cells. We considered two independent replicates for Parse and 10x platforms, which were derived from the same healthy individuals, H1 and H2. For Parse data, we multiplexed H1 and H2 with another nine human PBMC samples. By doing so, we maximized the capacity of the kit and allowed demultiplexed sample comparison with 10X. In general, both platforms produced high-quality data with desired number of cells, genes, and cell type clusters recovered. Overall, the data quality was largely consistent across the two biological replicates within each platform. However, we compared the relative strengths and weaknesses of each platform and assessed the impact of key features for most scRNA-seq studies, including library efficiency, sensitivity in gene detection, potential biases in gene quantification, and the capability to identify cell types accurately.

We selected peripheral blood mononuclear cells for our platform benchmarking since they are relatively easy to isolate, and the number of viable cells sequenced is typically high (>70%) after freezing. These factors minimize potential bias introduced during sample collection and preprocessing, allowing us to focus on the platform comparison. Moreover, PBMCs serve as a good reference cell system for our comparative analysis since they have been extensively studied with regards to their cell type composition and gene expression patterns [11, 15-16].

When compared with the 10x platform, Parse exhibits a two-folds decrease in cell recovery rate and in ∼13% fewer valid reads. This difference in library efficiency can be attributed to the distinct designs of the two platforms. In the 10x platform, a single droplet in the microfluidic channels aims to encapsulate one bead and a single cell. Cell lysis, barcoding, and reverse transcription all occur within this droplet. The number of cells in a droplet follows a super-Poissonian distribution, resulting in ∼80% bead occupancy and a cell recovery rate of about 50% [16]. On the other hand, Parse does not physically partition cells; instead, it fixes and permeabilizes all cells in a single mixture. Barcoding and reverse transcription are accomplished through four rounds of cell splitting and pooling in a multi-well plate. While this iterative process enhances sample throughput, it may compromise efficiency of library preparations. The repeated transfer of cells between wells and tubes increases the likelihood of cell loss during the procedure. The mechanical forces involved in this process can result in a higher rate of cell breakage, leading to the removal of these cells as background noise during the washing steps. These factors may explain why in Parse, approximately twice the number of cells are needed to be loaded in order to obtain a comparable number of cells recovered by the 10x system. Despite the lower cell recovery rate in libraries, Parse had fewer cells discarded in the quality control step, which can be attributed, for the greatest part, to a lower multiplet rate in Parse platform compared with 10x. 10x employs a single reverse transcription step to attach all barcodes, whereas Parse involves one reverse transcription step and three subsequent ligation steps. The inclusion of these four sequential reactions in the Parse can increase the likelihood of barcodes failing to attach to the cDNA molecules. This disparity between single and sequential barcoding could explain why Parse exhibits a relatively higher fraction of reads without valid barcodes. Consequently, these invalid reads occupy space in the sequencing library, therefore requiring an increased sequencing depth for Parse libraries in order to ensure optimal transcriptome coverage and accuracy of gene expression quantification.

Our analysis revealed a significantly higher proportion of ribosomal protein coding genes in the 10x data when compared with Parse. Notably, these findings are not unique to our study, as similar observations have been reported between SMART-seq and 10x platforms, whereby 10x data was enriched (2.6-7.2 fold) for ribosomal transcripts [28]. We expanded this comparative analysis to include a publicly available PBMC dataset generated using 12 distinct scRNA-seq protocols/platforms [20]. The proportion of ribosomal protein coding genes was the highest in 10x, though other non-droplet based platforms also showed a relatively high proportion (Supp. Figure 4G). The reasons behind this platform-specific bias in ribosomal protein coding gene enrichment require further investigations.

Gene detection rates can be increased by achieving greater sequencing depths. To compare the gene detection sensitivity of the two platforms in an unbiased way, the sequencing depth was standardized to 20,000 reads per cell for each sample. Our observations revealed that Parse detected approximately 1.2-fold more genes than 10x in T cells, NK cells, and B cells, but not in monocytes. This difference can be attributed to the fact that the sensitivity of the 10x platform relies on the mRNA quantities present in cells [20]. The relatively low mRNA quantities of T cells, NK cells, and B cells [29] affect the performance of 10x in detecting lowly-expressed genes. Conversely, monocytes provide higher RNA quantities [29] for 10x to start with, thereby enabling enhanced gene detection performance.

Higher gene detection sensitivity can greatly benefit downstream analysis of scRNA-seq data [14]. With Parse we observed higher clustering performance and greater power in distinguishing cell types with cell type-specific gene signatures. However, known cell type marker genes were lowly expressed in Parse data compared to 10x. This may be attributed to Parse’s tendency to detect longer and low GC content genes, as opposed to 10x. Therefore, when annotating cell types in Parse data, an alternative approach such as mapping reference datasets with known cell type information [30] may be more suitable than relying solely on a few specific marker genes.

Despite some differences between the two platforms, our analyses suggest that the scRNA-seq data demultiplexed from Parse have comparable quality to those obtained by a single 10x library. We provide a concise high-level summary of the relative strengths and weaknesses of each platform in Table 1. While it is not our aim to advise the use of a specific platform, we believe these analyses and findings can be helpful to facilitate decision-making processes and study design for scRNA-seq studies that require multiplexing.

## Methods

### Peripheral blood mononuclear cells (PBMCs) collection

PBMCs were isolated from EDTA anticoagulated venous blood from donors using a FicollR Paque Plus (Sigma Aldrich, # GE17-1440-02) solution, following standard density gradient centrifugation methods. The collected PBMC layers were then subjected to red blood cell lysis using ACK lysis buffer (Thermo Fisher, # A1049201), and then the PBMCs were suspended in a freezing media composed of 90% fetal bovine serum (FBS, HyClone) and 10% dimethyl sulfoxide (DMSO) and frozen at -80°C for future use.

### Single-cell suspension preparation

Frozen vials of PBMCs were rapidly thawed in a 37°C water bath until pea-sized frozen cryopreserved cells remained. Thawed PBMCs were quenched by adding 12 mL 37°C pre-warmed human serum (Sigma Aldrich, # H4522) and were centrifuged at 500 x g for 5 min at room temperature. The cell pellet was resuspended in 0.5 mL of pre-warmed human serum and rested in the 37°C, 5% CO2 incubator for 30 mins. After resting, cell solutions were passed through a 40 mm Falcon cell strainer (Thermo Fisher, # 08-771-1), and then centrifuged. Single live PBMCs were selected using fluorescence-activated cell sorting (FACS) with gating based on forward scatter (FSC), side scatter (SSC), and DAPI dye (Thermo Fisher, #D1306). The purified PBMCs were then resuspended with 1 mL 1X PBS containing 1% bovine serum albumin (BSA, Sigma Aldrich, # B2518).

For the 10x Genomics platform, the cell count for each sample was determined using the TC20™ Automated Cell Counter (Sigma Aldrich). Approximately 16,000 cells per sample were used for library construction, utilizing the Chromium Next GEM Automated Single Cell 3ʹ Reagent Kits (10x Genomics). For the Parse Biosciences platform, the cells were prepared using the Fixation Kit (Parse Biosciences) according to the manufacturer’s protocol. After fixation, the cell count was determined using the TC20™ Automated Cell Counter (Sigma Aldrich). A total of 390,000 cells from 11 donors was used, resulting in an average of approximately 35,500 cells per donor. These cells were subjected to the Single Cell Whole Transcriptome Kit v2 (Parse Biosciences) for library construction. The scRNA-seq libraries from 10x Genomics and Parse Biosciences platforms were sequenced using the Illumina NovaSeq 6000 platform, generating paired-end 200-bp reads as follows: for 10x, H1: 203,134,663 and H2: 254,103, 388; for Parse, H1: 297,237,012 and H2: 262,086,443 (Supp. Table 1).

### Parse and 10x data downsampling

For Parse data, fastq files from 8 sub-libraries were demultiplexed into 11 samples using *split-pipe*pipeline (v1.0.4p) from the Parse Biosciences (https://www.biorxiv.org/content/10.1101/2022.08.27.505512v1.full.pdf). Demultiplexed fastq files from sample H1 and H2 were used for downsampling. For 10x data, each sample is prepared and sequenced in a single library. We directly used these two fastq files for downsampling. Read downsampling was achieved for each sample by using *seqtk* (v1.3-r106) with command ‘*seqtk sample*’. Random seed was fixed at 100 with argument *-s*.

### Parse and 10x data preprocessing and quality control

The same human reference genome GRCh38/hg38 was used to map and quantify gene expression for 10x and Parse data. Human genome reference files are downloaded from Ensembl database (https://ftp.ensembl.org/pub/release-108/fasta/homo_sapiens/dna/Homo_sapiens.GRCh38.dna.primary_assembly.fa.gz; https://ftp.ensembl.org/pub/release-108/gtf/homo_sapiens/Homo_sapiens.GRCh38.108.gtf.gz).For 10x data, read mapping and gene expression quantification were done using *CellRanger* (v7.1.0) [16] using default parameters except for *--include-introns=true*. For Parse data, *split-pipe* (v1.0.4p) was used with the following parameters: *--mode all --chemistry v2 --kit WT --kit_score_skip --no_allwell*.

Gene expression matrices were loaded into the R package *Seurat* (4.3.0) [31] for quality control and downstream analyses. We removed cells fulfilling any of these criteria: (1) number of UMIs ≤ 1500, (2) number of UMIs ≥ 20,000, (3) number of genes ≤ 500, and (4) percentage of mitochondrial genes ≥ 15%. Cell doublets were detected and removed using R package *DoubletFinder* (2.0.3) [32]. Genes expressed in less than 8 cells are discarded. Gene counts are then normalized and transformed with *NormalizeData()*, which normalizes the total counts for each cell by the total counts, multiplies this by a scale factor 10,000 and log-transforms the result.

### Cell type annotation

Cell type annotation was done separately for Parse and 10x data using the same procedure as follows. First, highly variable genes (HVGs) were obtained using the *FindVariableFeatures()* function with default parameters. The top 2000 HVGs, which were decreasingly ordered based on dispersion, were selected as input to downstream analysis. Data were then scaled using *ScaleData()* function with parameter *vars.to.regress = c("nFeature_RNA", "percent.mt")*. Dimension reduction was performed using *RunPCA()* and *RunUMAP(dims=1:20)* functions. The top 20 principal components (PCs) were used to construct nearest-neighbor graphs, and identify cell clusters using the *FindNeighbors()* and *FindClusters()* functions of the *Seurat* (4.3.0) R package.

Cell type identity was confirmed using (1) consensus annotation from manual annotation with markers obtained from [14, 33, 34] and (2) reference-based annotation with *SingleR* (2.0.0) R package [35]. For manual annotation, 12 major cell types are identified, as follows: CD4+ T cells (CD3D+CD4+), CD8+ T cells (CD3D+CD8A+), MAIT cells (SLC4A10+), NK cells (NKG7+NCAM1+), CD14+ Monocytes (LYZ+CD14+), CD16+ Monocytes (LYZ+FCGR3A+), cDC (FCER1A+), pDC (CLEC4C+), B cells (MS4A1+), Plasmablasts (MZB1+), Proliferating cells (MKI67+) and HSPC (CD34+) (Supp. Table 4). For reference-based annotation, *SingleR()* function was used to calculate the gene expression correlation with the reference dataset from Monaco et al. [29] using default parameters.

A consensus cell type annotation between the manual and reference-based annotations was obtained using following steps: (1) compare manual and *SingleR* annotations, if they are identical, leave the cell type annotation as it is; (2) if one of the two annotations is at higher resolution (i.e. a cell type (e.g., CD4+ T cell) is a subset of another cell type (e.g., T cell)), use the annotation with the higher resolution; (3) if the two annotations are still inconsistent, use canonical markers to re-annotate the cell; (4) remove cells that express conflicting cell type markers (e.g. cells co-expressing CD3D+ T cell and MS4A1+ B cell markers) as potential doublets.

### Library efficiency

The number of reads without valid barcodes were obtained and summarized by *split-pipe* (v1.0.4p) and *CellRanger* (7.1.0) separately, and divided by total number of reads to obtain the fraction of valid reads for each sample. *QualiMap* (2.2.2) [36] was used to quantify reads uniquely mapped to exons, introns, intergenic and to overlapping-exon regions between any two overlapping genes. For each sample, the cell capture efficiency was calculated by dividing the number of cells output from *split-pipe* (v1.0.4p) and *CellRanger* (7.1.0) by the total number of loaded cells.

### Dropout probabilities estimation

For each sample and separately for each dataset generated by 10x or Parse, we estimated the dropout probabilities on the down-sampled data (20,000 reads/cell). We randomly sampled 50 cells for each sample to avoid bias from the different number of total cells in each sample. The estimation was done with *scde* (1.2.1) R package [37], which models the gene expression matrix (UMI counts) as a mixture of two probabilistic processes: one is a negative binomial process which accounts for amplified mRNA, the other is the poisson process to account for dropout mRNA. Dropout probabilities for each cell were calculated with *scde.failure.probability()* function with default parameters. Average dropout probabilities were summarized and plotted against gene expression magnitudes (UMI counts).

### Correlation analysis

For each sample we randomly sampled 7,000 cells from the down-sampled data (20,000 reads/cell) which were used for gene expression correlation analysis. Only genes expressed in all samples were included in this analysis. Gene counts of commonly expressed genes were averaged across all cells in each sample. These average gene counts were used to assess the Pearson’s correlation between Parse and 10x in each sample H1 and H2 separately. Pearson’s correlation coefficients were calculated using the *cor()* function in base R with ‘*method = ‘pearson*’.

### Platform-specific marker genes

Differentially expressed genes (DEGs) between Parse and 10x data were identified using *FindAllMarkers()* function from *Seurat* (4.3.0) R package with parameters: *only.pos = TRUE, logfc.threshold = 0.25, min.pct = 0.1, test.use = ’MAST’, latent.vars = c(’nFeature_RNA’)*. Up-regulated DEGs with average log2 fold change larger than 1 and adjusted p value smaller than 0.01 in Parse and 10x, respectively, were identified as marker genes of each platform. The length and GC content for each gene were downloaded from BioMart (http://asia.ensembl.org/info/data/biomart/index.html).

### Ribosomal protein coding genes

Ribosomal protein coding genes were identified using gene symbols starting with RPS/RPL. Single cell RNA-seq data generated with 12 different methods [20] were downloaded from https://github.com/elimereu/matchSCore2/tree/master/data.

### Silhouette score

Down-sampled data (20,000 reads/cell) was clustered using graph-based clustering with first 10 PCs at resolution of 0.4, 0.5, 0.6, 0.7, 0.8, 0.9, 1.0, 1.1, 1.2, 1.3, 1.4, 1.5 with functions *FindNeighbors()* and *FindClusters()* in *Seurat* (4.3.0) R package. We used *silhouette()* function from *cluster* (2.1.3) R package to compute Average Silhouette Width (ASW) scores based on Euclidean distance calculated by *dist()* function in *stats* (4.2.2) R package.

### Cell type signature score

Bulk RNA-seq data from sorted immune cells [27] are obtained by calling *DatabaseImmuneCellExpressionData()* function from *celldex* (1.4.0) R package [35]. Differential expression analysis was carried out for each cell type’s RNA-seq data using *findMarkers()* function from *scran* (1.22.1) R package [38]. Up-regulated genes with FDR less than or equal to 0.01 were used as cell type gene signatures. *AUCell_calcAUC()* from the *AUCell* (1.16.0) R package [39] was used to calculate the AUC of each cell type gene signature in each cell. Median value of AUC scores across all the cells in that cell type were summarized and indicated in violin plots as the measurement of dataset’s capability in cell type annotation.

---

**Supplementary Figure 1.**
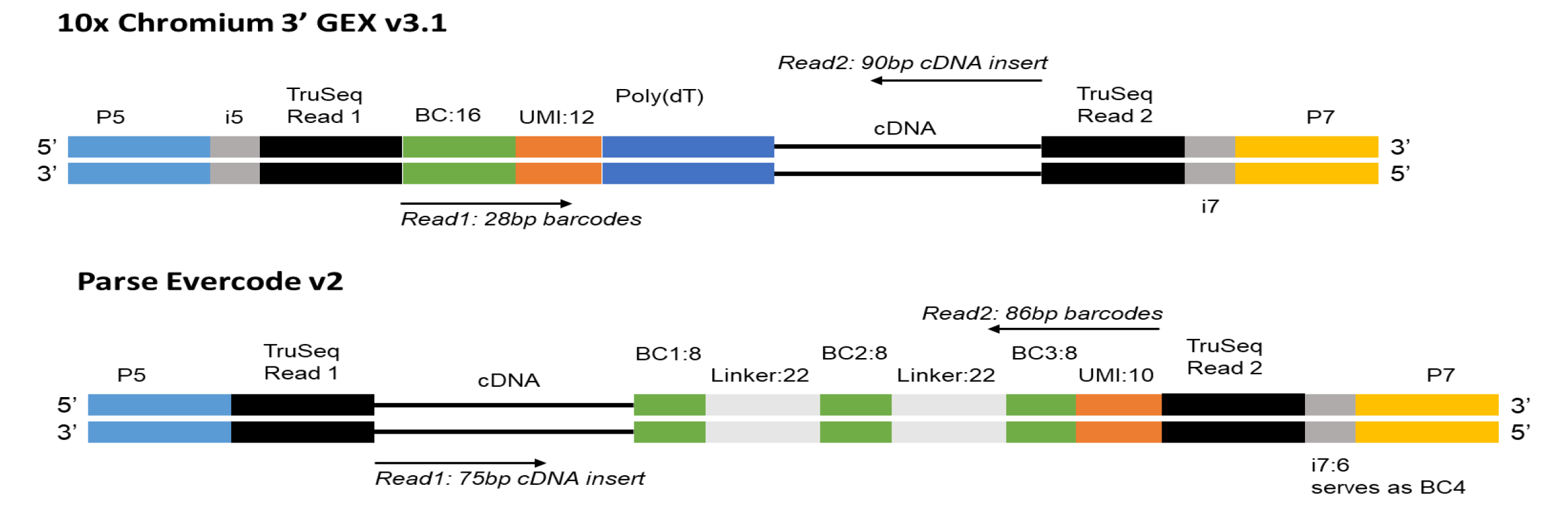
cDNA library structure. BC: barcode, UMI: unique molecular identifiers.

**Supplementary Figure 2.**
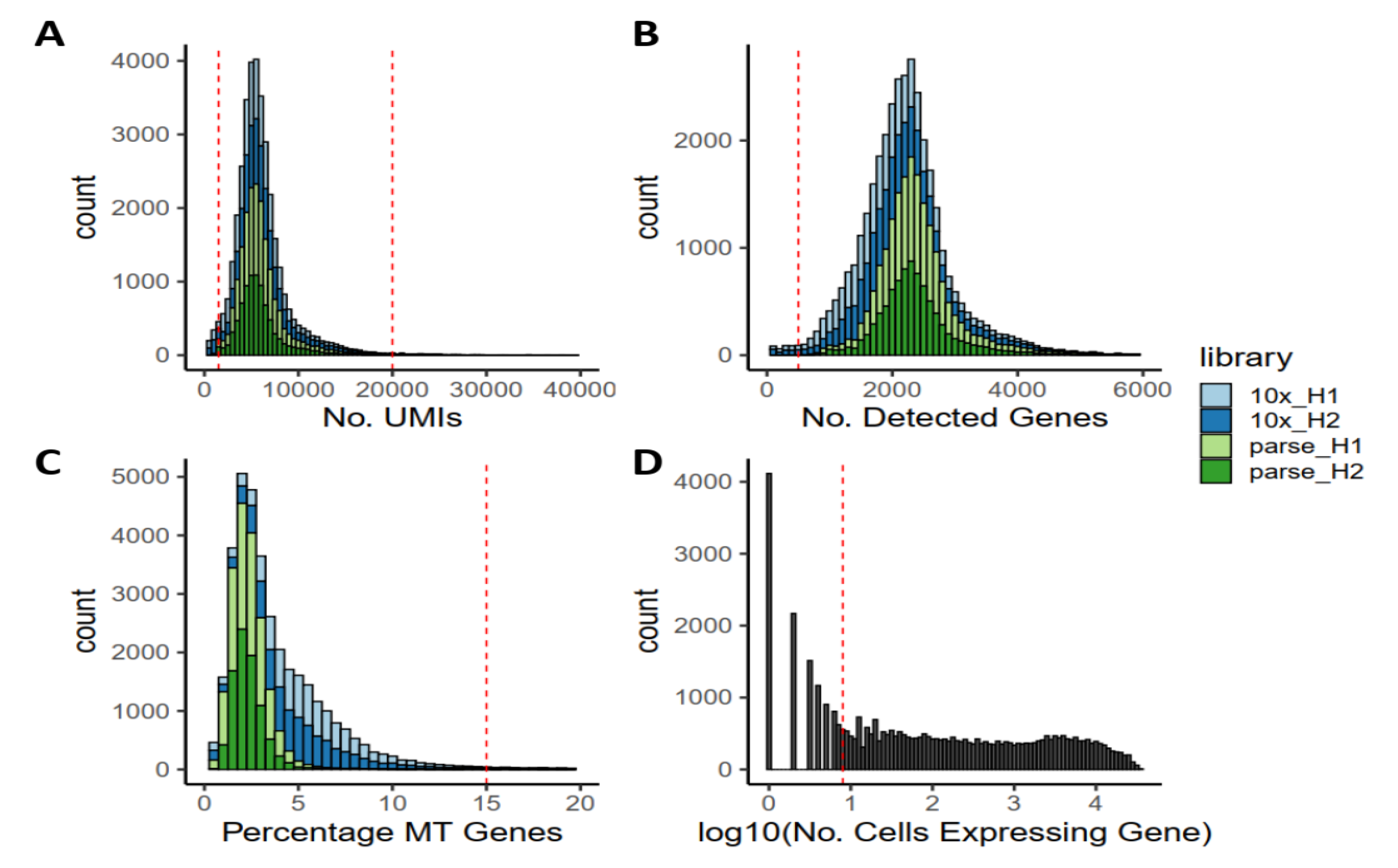
Quality control. **A)** Distribution of UMIs detected per cell. **B)** Distribution of genes detected per cell. **C)** Distribution of percentage of UMIs mapped to mitochondrial genes per cell. **D)** Distribution of number of cells expressing the gene. Red dashed lines represent thresholds we used to filter low-quality cells and lowly-expressed genes.

**Supplementary Figure 3.**
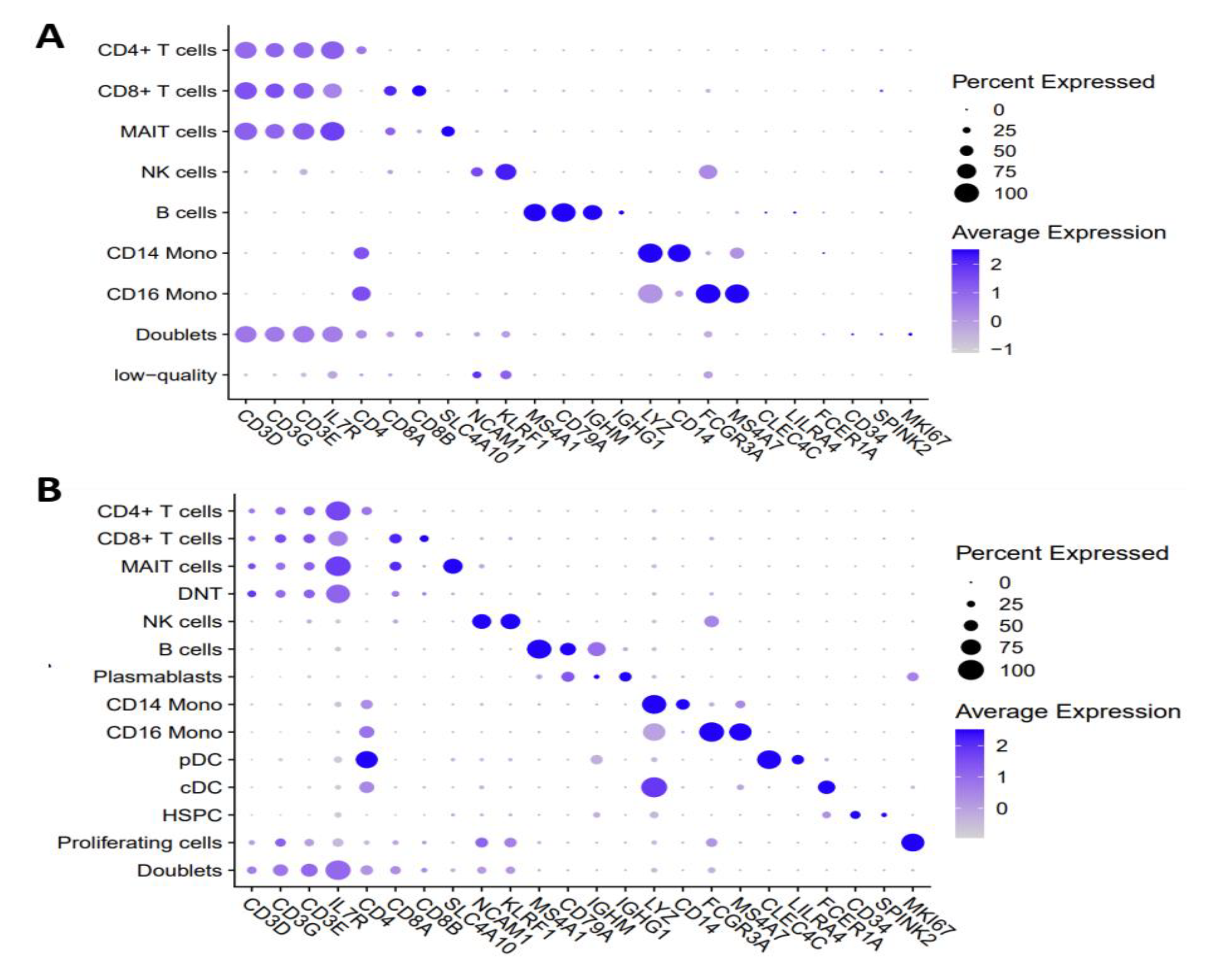
Cell type annotation. **A)** Dotplot of average expression of cell type marker genes in 10x data. **B)** Dotplot of average expression of cell type marker genes in Parse data.

**Supplementary Figure 4.**
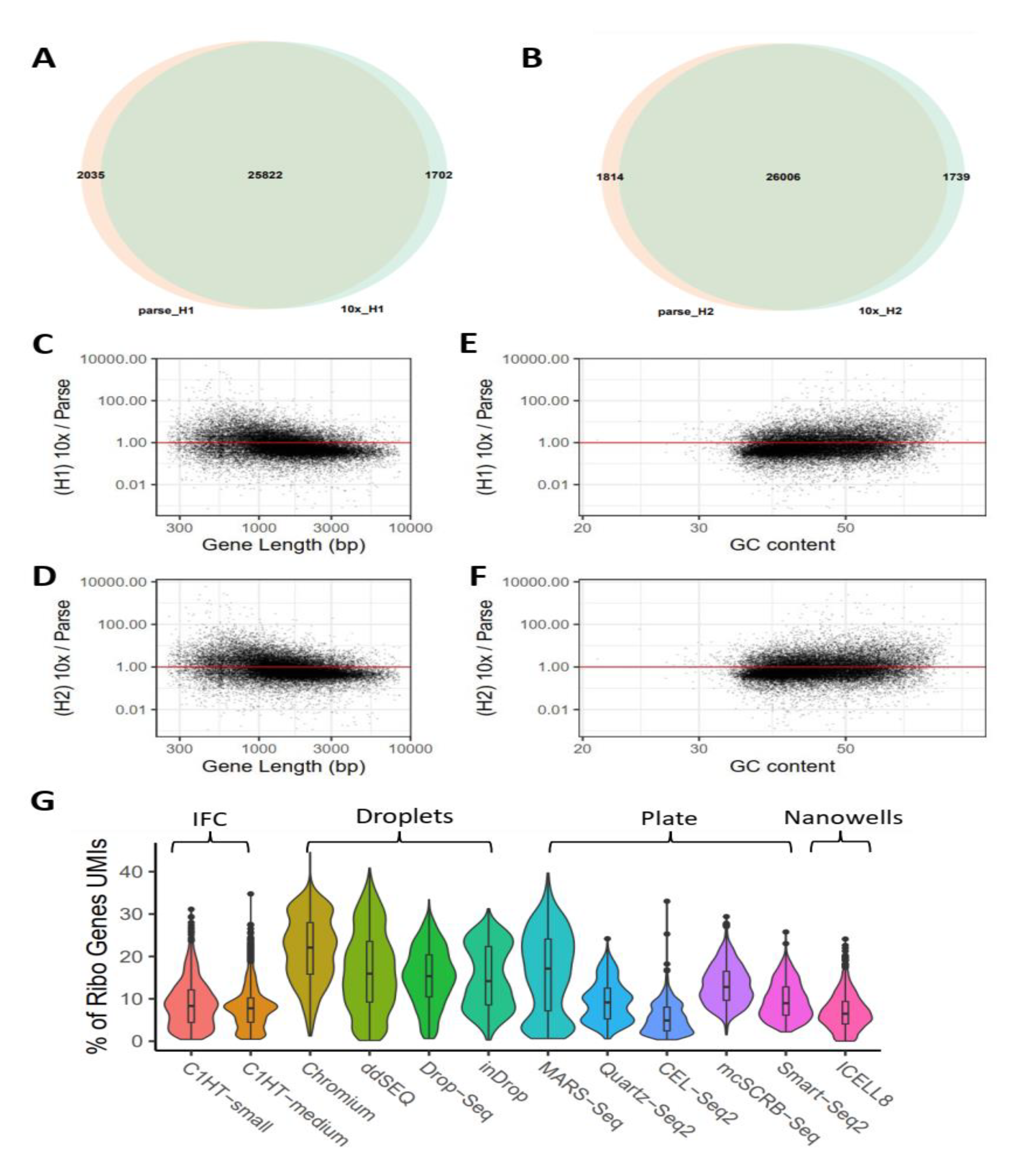
Differences in gene quantification. **A, B)** Number of shared and unique genes in sample H1 (A) and H2 (B) between Parse and 10x. **C, D)** Scatterplots of gene expression ratio and gene length in sample H1 (C) and H2 (D). Each dot represents one gene that is expressed in all samples. Y axis is the ratio of average gene expression between 10x and Parse. **E, F)** Scatterplots of gene expression ratio and GC content in sample H1 (E) and H2 (F). Each dot represents one gene that is expressed in all samples. Y axis is the ratio of average gene expression between 10x and Parse. **G)** Ribosomal protein coding gene abundance in human PBMC datasets generated with 12 different scRNA-seq protocols. Cell capture technology of each protocol is indicated (IFC: integrated fluidic circuit; C1HT: Fluidigm; Chromium: 10x Genomics).

**Supplementary Figure 5.**
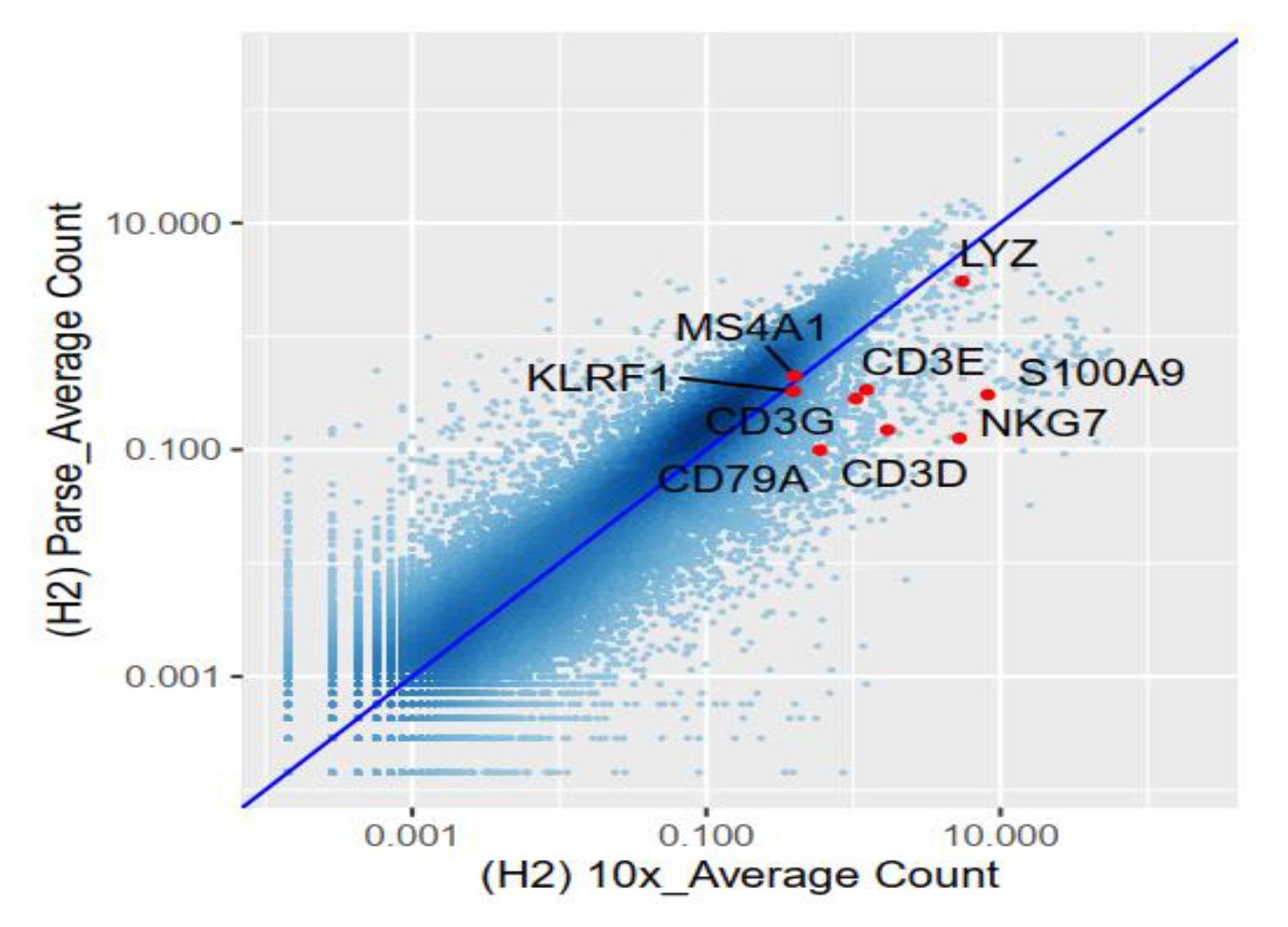
Gene expression correlation. Scatterplot showing gene expression correlation between two platforms in sample H2 colored by density. X-axis is the average UMI count for each gene in 10x data and Y -axis is the average UMI count for each gene in Parse data. Cell type markers are highlighted with red dots.

**Supplementary Table 1.**
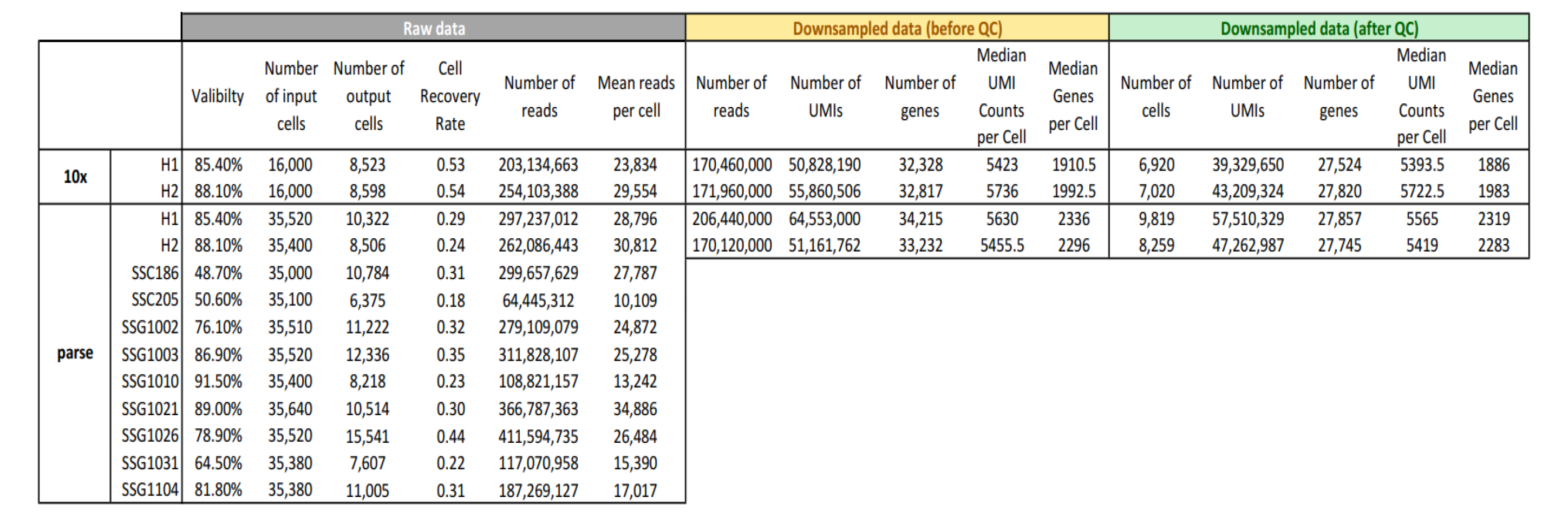
Overview of PBMCs samples in this study

**Supplementary Table 2.**
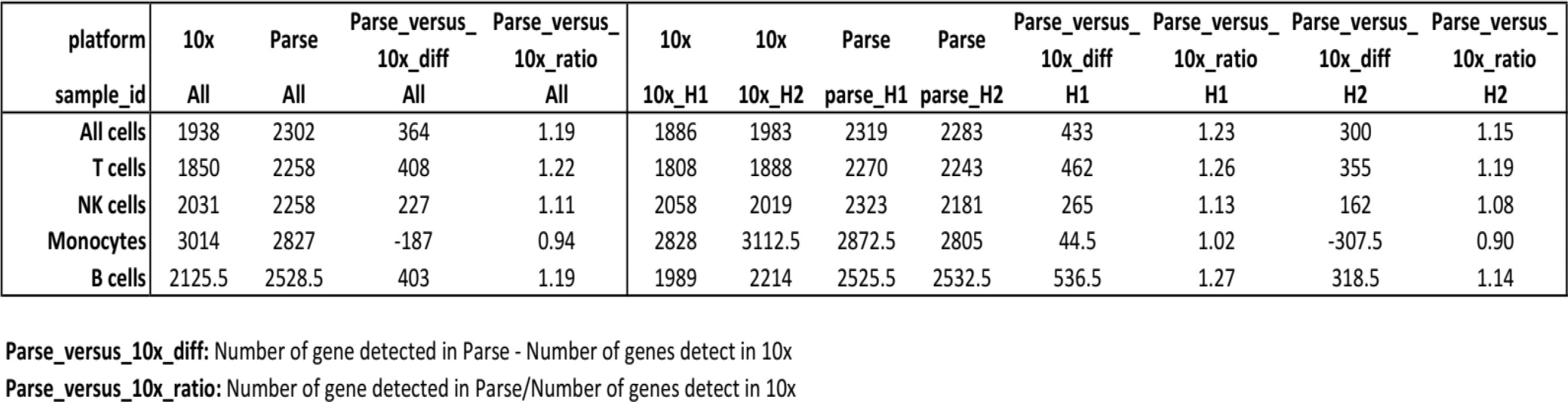
Median number of genes detected in each protocol/sample

**Supplementary Table 3.**
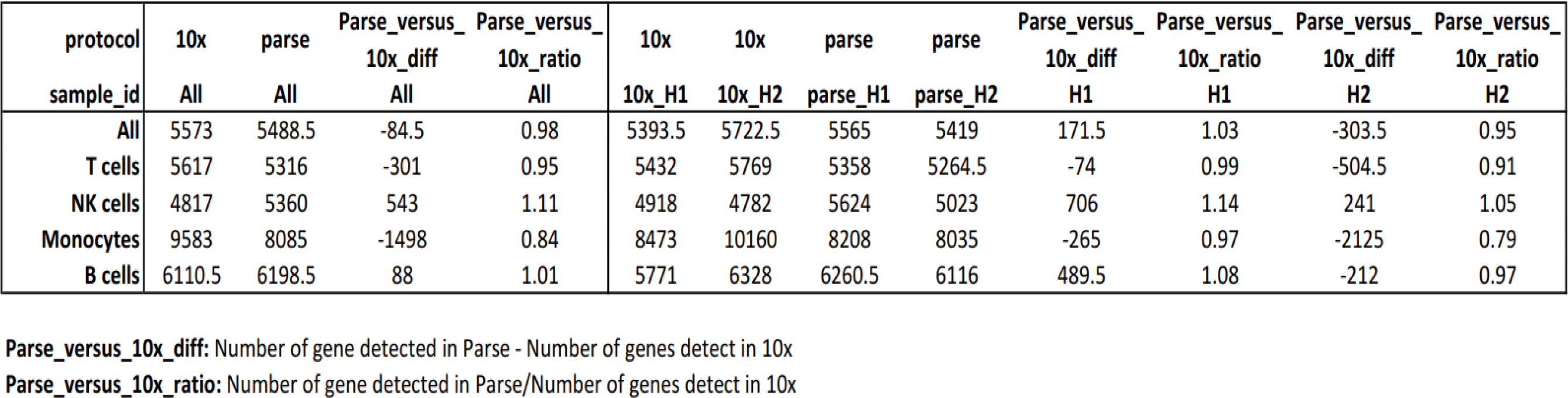
Median number of UMIs detected in each protocol/sample

**Supplementary Table 4.**
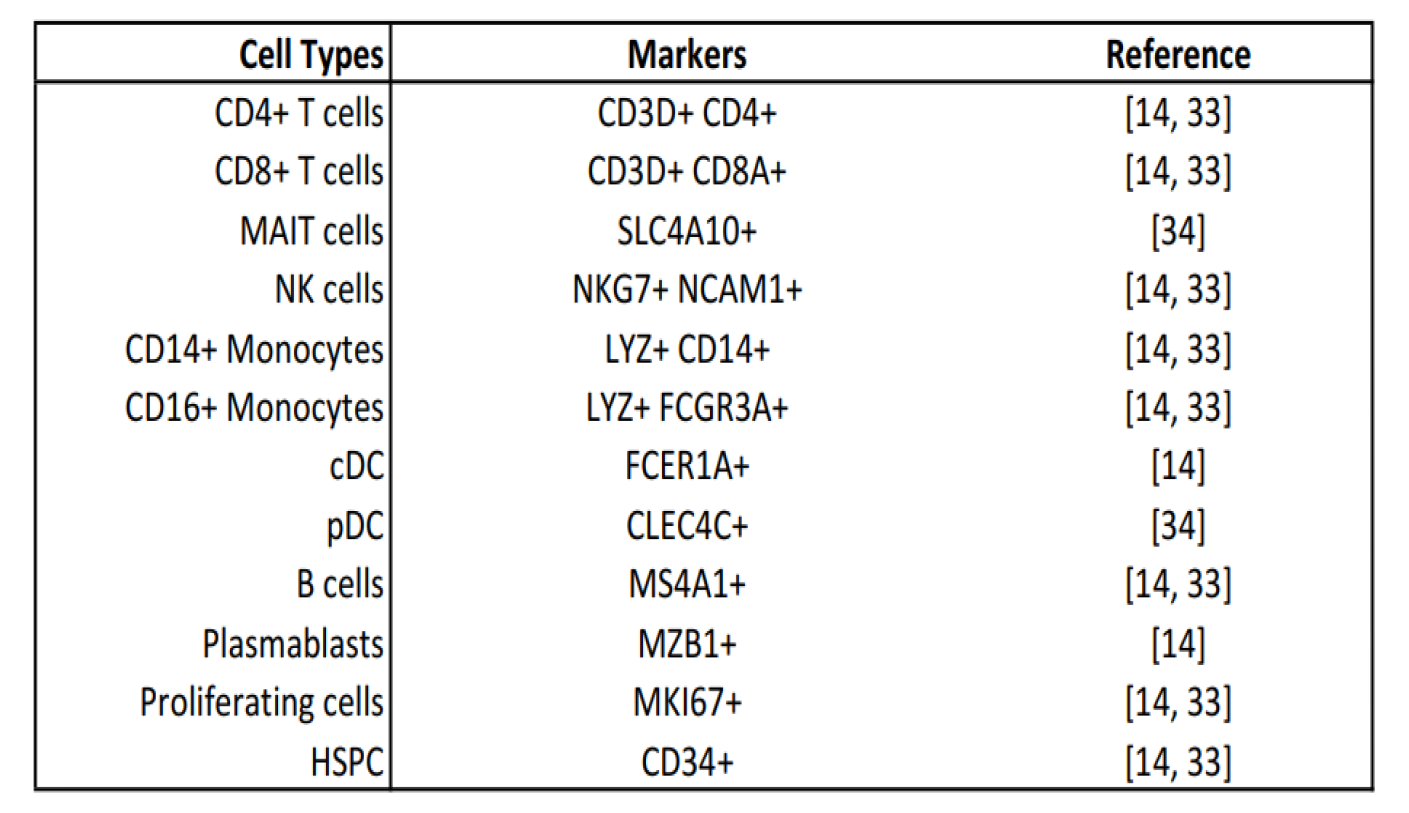
Canonical markers used for cell type annotation

